# Skin specimen as an alternative to brain tissue for post mortem rabies diagnosis in animals to scale up animal rabies surveillance

**DOI:** 10.1101/2021.09.01.458510

**Authors:** Sujatha Aparna, Abraham Susan Swapna, GS Ajithkumar, Sujatha Chintha, PR Prathiush, S Nandakumar, Joseph Baby

**Author notes:** Corresponding author: Dr Sujatha Aparna, MVSc, Veterinary Surgeon, State Institute for Animal diseases, Palode, Government of Kerala, India. Contact number: 09447958081. **Author contributions: SA:** Conceptualization, Methodology, Investigation, Formal analysis, Original draft preparation, Review and editing, Project administration; **ASS:** Methodology, Resources, Investigation, Formal analysis, Original draft preparation, Review and editing; **GSA:** Methodology, Investigation, Review and editing; **SC:** Formal analysis, Original draft preparation, Review and editing; **PRP:** Investigation, Review and editing; **SN:** Review and editing; **JB:** Project administration, Review and editing.

## Abstract

**Background:** Detection of the virus or some of its specific components using WHO and OIE recommended standard laboratory tests is the only way to get a reliable diagnosis of rabies. Brain tissue is the preferred specimen for post-mortem diagnosis of rabies in both humans and animals. Higher biosecurity requirements, skill and transportation facilities required for collection and transport of brain or whole carcass to the laboratory is one of the reasons for the poor rabies surveillance in animals. Point of care testing with simple, reliable and easy to operate devices would be an ideal approach for providing rapid results.

**Methods:** The study evaluated diagnostic performance of two reference tests, DFAT and RTPCR on skin specimen, to assess its suitability as an alternative of brain tissue for post mortem rabies diagnosis in animals. Brain tissue and skin sample belonging to different species of animals (n=90) collected at necropsy were compared using Fluorescent Antibody Test and RT PCR, internationally approved methods for rabies diagnosis.

**Results:** Validation of RT-PCR on skin and DFAT on skin in comparison with DFAT on brain as gold standard gave a sensitivity of 98% (95% CI:94.1-100) and 80% (95% CI:71.8-88.2) respectively. Specificity was 100% in both tests.

**Conclusion:** The findings highlight the potential of skin specimen for improving rabies surveillance in animals especially in resource poor countries.

**Author Summary:** The present study was undertaken to define a reliable protocol for diagnosing animal rabies using skin specimen, a superficial tissue sample which is collected in a non-invasive manner for post-mortem diagnosis. Our aim was to design a protocol to replace the classical post-mortem diagnostic method that uses brain biopsy with an ultimate target of stepping up rabies surveillance in animals. Brain tissue and skin sample belonging to different species of animals collected at necropsy were compared using Fluorescent Antibody Test and RT PCR, internationally approved methods for rabies diagnosis. The study established that RT PCR on skin specimen is rapid, sensitive and specific, opening its potential as an ideal rabies surveillance tool overcoming the logistical challenges of carcass transportation to reference laboratories and alleviating biosafety concerns associated with brain collection. The study highlights the potential of skin specimen for improving rabies surveillance in animals especially in resource poor countries.

## Introduction

Rabies is still a significant public health problem in many parts of the globe. Most cases of human rabies occur in Asia and Africa with majority of cases transmitted through dog bites. Elimination of human rabies is feasible and sustainable if rabies is controlled at its source. There is a need to scale up dog rabies surveillance and reporting in developing countries for the global dream of canine rabies elimination to become a reality.

India contributes about 40% of the global disease burden with an annual estimated 20,000 deaths.(1) However there is a dearth of data relating to disease burden and distribution in animals. This is mainly due to insufficient laboratory capacity at the field level and poor sample flow to reference laboratories. Various factors are responsible for the underestimation of animal rabies deaths. Post mortem brain testing, with sensitivity nearly 100%, is the gold standard method of rabies diagnosis.(2,3) Maintaining cold chain during transport, bio-security issues encountered in transporting whole carcass, decapitation of head and trouble of opening skull in field conditions discourage veterinary practitioners from reporting and confirming rabies suspected cases. Collection of brain through foramen magnum though widely propagated as a field sampling method requires certain amount of skill and expertise. Hence there is need for a non- invasive alternative sample which can be collected easily at the same time demonstrating equal efficiency as cerebral biopsy.

The present study was undertaken to define a reliable protocol for diagnosing animal rabies using skin specimen, a superficial tissue sample which is collected in a non-invasive manner for post mortem diagnosis.(4,5) Our aim was to design a protocol to replace the classical post mortem diagnostic method that uses brain biopsy with an ultimate target of stepping up rabies surveillance in animals.

## Materials and Methods

### Study design and Setting

This study was planned as a diagnostic test evaluation. It was done at State Institute for Animal Diseases (SAID) Palode, the State referral laboratory for animal disease diagnosis in Kerala, India where samples from all parts of the state are received for rabies diagnosis. On an average two hundred samples are received annually for rabies diagnosis. The study period was from June 2020 to January 2021.

### Study material

The study material consisted of rabies suspected animal carcasses presented to the State Institute for Animal Diseases through routine rabies surveillance program of State department of Animal Husbandry. The sample size was calculated using the formula for estimation of proportion (n= zα^2^pq/d^2^). To estimate a sensitivity of 82% as per a previous study of *Yale etal*.,2016(6) at 95% confidence level and 10% absolute precision, 56 positive samples were needed. With the assumption that 60% of the samples received will be positive for rabies a total sample size of 90 was taken. Consecutive samples received were included in this study. A total of 90 animals (n) were tested which included 86 domestic dogs, 2 wild dogs, 1 buffalo and 1 rabbit.(3) Carcasses in advanced putrefactive stage that preclude the identification of anatomical sites were excluded from the study.

Brain tissue was collected by opening skull and pooled material from specific anatomical sites like stem, cerebellum and hippocampus was used as the sample. All samples were tested by DFAT on the same day. A full length skin sample taken from the nape of neck including a portion of subcutaneous tissue and muscle from each case was subjected to DFAT by a different investigator. RT PCR was conducted on skin and brain specimen in batches. Samples were stored at −20°C until testing. DFAT results were blinded from the investigator who conducted RT PCR.

### Test methodology

#### Direct Fluorescent Antibody Test (DFAT)

DFAT was carried out according to Meslin *et al.,* 1996.(7) Impression smear preparations of the brain samples were air dried and placed in a Coplin jar containing chilled acetone and fixed at 4°C for one hour. The slides were air dried and incubated with anti-rabies nucleocapsid conjugate (Bio-Rad, France) for 35 min at 37°C in a humid chamber and further washed with phosphate buffered saline (PBS) of pH 7.2 in two successive washes for 5-10 min, air-dried and mounted with buffered glycerol and then visualized under an fluorescent microscope (Olympus BX51) at 400X magnification. Results as bright/dull/dim apple-green round to oval intracellular fluorescence accumulations or no fluorescence were graded as per *Tepsumethanon et al.,* 1997.(8)

Touch impression smears taken from skin triturate was also subjected to DFAT using same protocol described above.

### Real time PCR (RT QPCR)

#### RNA extraction

Samples were processed by the use of RNeasy Mini Kit (Qiagen), as per the manufacturer’s recommended protocol. Concentrations of DNA were calculated by the use of a spectrophotometer.

### Real time PCR (qPCR) analysis

#### Primersequences

1) Pan-lyssavirus-specific primers (HPLC purified)
a) JW12 RT/PCR primer 5’-ATG-TAA-CAC-CYC-TAC-AAT-G-3’
b) N165-146 PCR primer 5’-GCA-GGG-TAY-TTR-TAC-TCA-TA-3’
2) Multispecies Beta Actin primers (HPLC purified) (Internal control)
a) BatRat Beta-actin forward primer 5’-CGA-TGA-AGA-TCA-AGA-TCA-TTG-3’
b) BatRat Beta-actin reverse primer 5’-AAG-CAT-TTG-CGG-TGG-AC-3’

#### Procedure

reaction mixture was prepared for both rabies primers and internal control primers separately as described below. The assay for β-actin must be positive in order to have confidence that RNA was isolated from the starting material:

| Reagent | Volume per reaction (micro litre) |
| --- | --- |
| Molecular grade water | 7.55 |
| Universal reaction mix containing a fluorescent stain (2×) SYBR Green | 10.0 |
| JW12 | 0.6 |
| N165-146 | 1.0 |
| RT enzyme mix | 0.25 |
| Total | 19.4 |
| Molecular grade water | 7.55 |
| Universal reaction mix containing a fluorescent stain (2×) SYBR Green | 10.0 |
| BatRat Beta-actin FP | 0.6 |
| BatRat Beta-actin RP | 1.0 |
| RT enzyme mix | 0.25 |
| Total | 19.4 |

#### Thermal cycler set up

**Thermal cycler set up:**
| Temperature ($^{\circ}$C) | Time | Cycles |
| --- | --- | --- |
| 50 | 10 minutes | 1 |
| 95 | 5 minutes | 1 |
| 95 | 10 seconds | 40 |
| 60 | 30 seconds |  |
| 95 | 1 minute | 80 |
| 55 | 1 minute |  |
| 55-95 | 10 seconds |  |

#### Interpretation

The specific fluorescence produced during the formation of amplicons/ PCR product from the genomic RNA (converted to cDNA) of Rabies virus was identified by the detector in the qPCR machine (CFX CONNECT, BIORAD). The quantitative result is expressed in terms of relative quantification. A negative result would not produce any detectable fluorescence in case of primers for rabies (JW12/ N165-146) and fluorescence in case of internal control BatRat Beta-actin only.

#### PCR controls

a sequenced gene product of specific primer (sample obtained from confirmed positive case of Rabies virus) was utilized as the positive control and negative samples for the disease were used as negative controls.

#### Data Analysis

Result of skin testing, the sample under evaluation, was compared against brain DFAT, the gold standard test for rabies diagnosis. Diagnostic performance of skin was estimated calculating sensitivity, specificity, positive and negative predictive values and accuracy using 2 × 2 table method. 95% confidence interval (95% CI) for the estimates obtained was determined.

## Results

Out of the 91 samples received during the study period, one was excluded from the analysis as the brain was autolysed beyond the identification of anatomical sites. Of the 90 remaining, 50 were tested positive and rest negative by DFAT on brain sample giving a percentage positivity of 55.5%. Animals positive on DFAT, reference method for rabies confirmation (OIE manual, 1998) in brain samples were considered as true positive animals (n=50) and the remaining (n=40) as true negative. RT PCR results on brain samples were 100% in accordance with DFAT results. Test positivity with DFAT and RT-PCR on brain is depicted in table 1.

**Table 1 :**
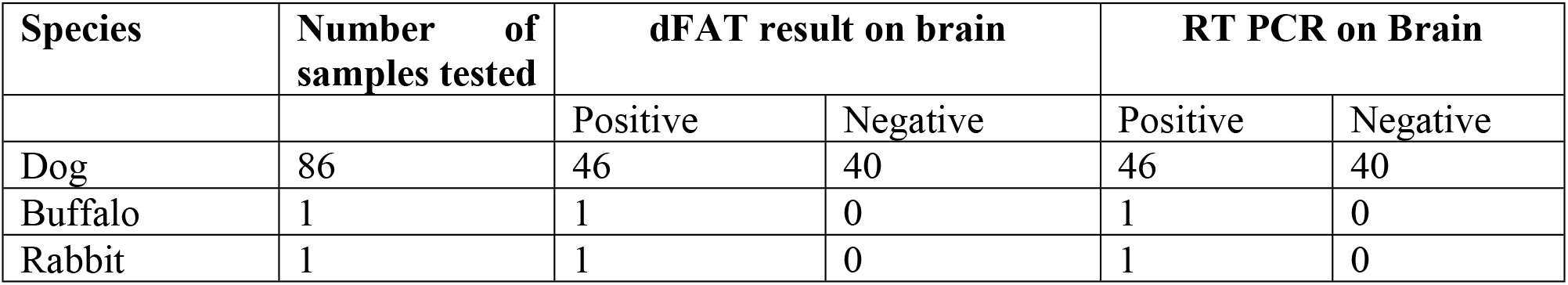
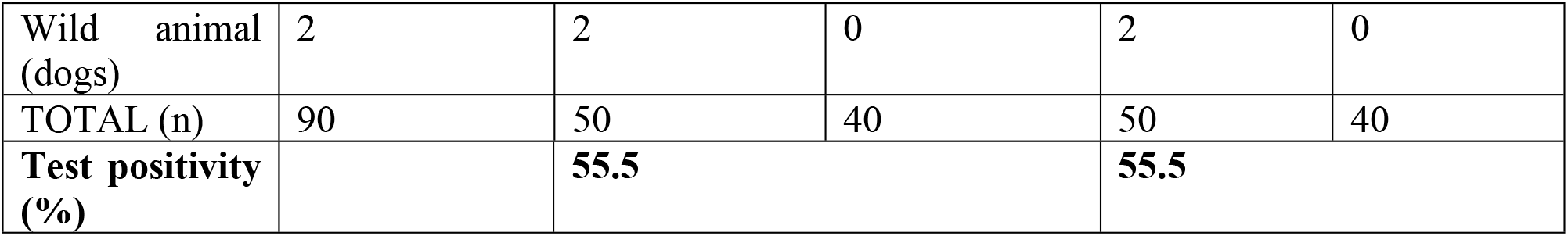
Detection of True Rabies Positive and True Rabies Negative Cases.

Of the 50 skin specimens of rabies positive animals tested, 49 were positive on RT PCR. None of the skin samples of rabies negative animals were positive in any of the two tests employed. However DFAT on skin impression smears could detect rabies only in 40 samples out of 50 true positive samples tested. No false positive reaction was observed on DFAT of skin specimens of negative animals. When brain testing gave test positivity of 55.5% (Table 1), skin testing demonstrated 44.4% and 54.4% respectively on DFAT and RT PCR (Table 2).

**Table 2:** Rabies positivity on skin specimen on various tests.

| Number of samples | dFAT on skin imprints | RT PCR on skin |
| --- | --- | --- |
| Positive : | 40 | 49 |
| Negative : | 50 | 41 |
| TOTAL (n) | 90 | 90 |
| Test positivity (%) | 44.4 | 54.4 |

Diagnostic performance of skin specimen using different reference techniques assessed by sensitivity, specificity, positive predictive value, negative predictive value and accuracy are depicted in table 3, 4 & 5. When sensitivity varied from 80% to 98% in two different reference tests, specificity was 100% in both tests. In the study, skin specimen when tested using molecular tests like RT PCR demonstrated 98% sensitivity revealing good potential as an alternative sample to brain tissue for rabies diagnosis in animals

**Table 3 :** Cross analysis of results of two specimens (skin vs brain) using DFAT.

|  |  | DFAT on brain (Gold standard) |  |  |
| --- | --- | --- | --- | --- |
|  |  | Positive | Negative | Total |
| DFAT on skin imprints | Positive | 40 | 0 | 40 |
|  | Negative | 10 | 40 | 50 |
|  | Total | 50 | 40 | 90(n) |

**Table : 4.** Cross analysis of results of two specimen (skin by RT PCR vs Brain by DFAT)

|  |  | DFAT on brain (Gold standard) |  |  |
| --- | --- | --- | --- | --- |
|  |  | Positive | Negative | Total |
| RT PCR on Skin | Positive | 49 | 0 | 49 |
|  | Negative | 1 | 40 | 41 |
|  | Total | 50 | 40 | 90 (n) |

**Table 5 :** Diagnostic performance of various tests on skin specimen for rabies diagnosis.

| Test | Sensitivity (95%CI) | Specificity | Positive Predictive Value | Negative Predictive Value(95%CI) | Accuracy (95%CI) |
| --- | --- | --- | --- | --- | --- |
| DFAT | 80<br>(71.8-88.2) | 100 | 100 | 80<br>(71.8-88.2) | 88.8<br>(82.4-95.2) |
| RT PCR | 98<br>(94.1-100) | 100 | 100 | 97.5<br>(94.4-100) | 98.8<br>(96.7-100) |

Grading of antigenic mass in skin and brain tissue using DFAT (Table 6 & Figure 1), revealed low antigen concentration in skin when compared to brain.

**Table :6.** Grading of antigen concentration by DFAT.

| Observation | Grade | Brain smear | Skin smears |
| --- | --- | --- | --- |
| Bright apple green fluorescence scattered throughout the field | 3+ | 46 | 1 |
| Dull apple green fluorescence scattered in min 50% of area | 2+ | 3 | 9 |
| Dim but visible fluorescence in min 10% of field | 1+ | 1 | 17 |
| TOTAL |  | 50 | 40 |

## Discussion

Animal rabies has not been regularly reported in resource poor developing countries. Testing on post mortem cerebral samples is the reference method for rabies diagnosis in animals. As sampling of brain tissue by opening the skull requires higher bio-security facilities, one need to transport whole carcass or decapitated head to reference laboratories which is often located at a distant place. There is a need for reliable non neuronal sample which can be easily collected and transported to the laboratory to scale up animal rabies surveillance.

Saliva and nuchal skin biopsy are proved to be diagnostic in animals.(9–12) Nuchal skin biopsy is reported to be more widely accepted sample for ante-mortem rabies diagnosis in humans and animals.(3,13–16) However sampling is difficult and risky in live rabies suspected animals and hence there are only limited studies evaluating its utility as an ante-mortem tool in animals(3,6,17) and only very few studies regarding its effectiveness as a post mortem sample.(6) Viral nucleocapsids are located in nerve endings surrounding base of hair follicles.(17) Skin is easy to collect, easily accessible in a non- invasive manner and hence got larger scope as a field sample. The result of the present study established that the window for post mortem rabies diagnosis with use of skin specimen is large and this sample effectively confirms the diagnosis of rabies. In our study the sample exhibited very high sensitivity and specificity (Table 1-6) in comparison to the gold standard specimen, the brain. The comparatively low sensitivity obtained DFAT could be due to the fact that the present study was conducted on skin imprints not in the standard method of frozen skin sections.(17,18) As expected viral load in skin sample was comparatively low in comparison to brain and might have been below the microscopic detection limit in the false negative cases detected in this study. The sensitivity of the test is likely to decline in cases of low antigenic load as it may happen in animals euthanized before the full clinical course. However, as the detection limits of molecular tests like RT PCR are very low, it would detect the infection even in cases of low antigenic mass.

The study established that RT PCR on skin specimen is rapid, sensitive and specific, opening its potential as an ideal rabies surveillance tool overcoming the logistical challenges of carcass transportation to reference laboratories and alleviating biosafety concerns associated with brain collection. To our knowledge, ours is the first study evaluating the utility of skin sample as a post mortem diagnostic tool involving different species of animals. Rabies incidence in wild animals is underestimated globally due to many factors. Lack of systematic surveillance tools, limited infrastructure for sample collection, difficulty in obtaining fresh samples and absence of trained skilled staff are cited among them. The study also suggests that skin could be useful in wild animal rabies surveillance as we could include two wild dogs in the study group and both of them were detected rabies positive by skin testing.

We recommend that suitability of skin as a post mortem tool is further evaluated on a larger sample set including wider species range. It is possible that different skin sites may show different level of sensitivity. There is evidence that muzzle skin sample of dog show higher sensitivity than nuchal skin due to high innervation. Autolysis is another challenge encountered in animal rabies surveillance especially in tropical countries due to high ambient temperature. It will be worthwhile to undertake studies to assess the effect of different factors like sampling time after death and skin collected from different areas to select the most promising site.

The study highlights the potential of skin specimen for improving rabies surveillance in animals especially in resource poor countries.

## Notes

### Competing Interest Statement

The authors have declared no competing interest.

## References

1. Sudarshan MK, Madhusudana SN, Mahendra BJ, Rao NSN, Ashwath Narayana DH, Abdul Rahman S, et al. Assessing the burden of human rabies in India: results of a national multi-center epidemiological survey. Int J Infect Dis. 2007 Jan;11(1):29–35.

2. Briggs D, Bourhy H., Cleaveland S., Cliquet F., Ertl H., Fayaz A., et al. WHO expert consultation on rabies. [cited 2021 Aug 30]; Available from: https://www.elibrary.ru/item.asp?id=13488298

3. Zieger U. Diagnosis of Rabies via RT-PCR on Skin Samples of Wild and Domestic Animals. Open J Vet Med. 2015;05(09):191.

4. Crepin P, Audry L, Rotivel Y, Gacoin A, Caroff C, Bourhy H. Intravitam Diagnosis of Human Rabies by PCR Using Saliva and Cerebrospinal Fluid. J Clin Microbiol. 1998 Apr 1;36(4):1117–21.

5. Warrell MJ, Looareesuwan S, Manatsathit S, White NJ, Phuapradit P, Vejjajiva A, et al. Rapid diagnosis of rabies and post-vaccinal encephalitides. Clin Exp Immunol. 1988 Feb;71(2):229–34.

6. Yale G, Mani RS. Utility of Skin Biopsy Sample for Rabies Diagnosis in Dogs. J Vet Sci Med Diagn [Internet]. 2016 [cited 2021 Aug 30];05(06). Available from: http://www.scitechnol.com/peer-review/utility-of-skin-biopsy-sample-for-rabies-diagnosis-in-dogs-uhB8.php?article_id=5643

7. Meslin F-X, Kaplan MM, Koprowski H, World Health Organization, editors. Laboratory techniques in rabies. 4th ed. Geneva: World Health Organization; 1996. 467 p.

8. Tepsumethanon V, Lumlertdacha B, Mitmoonpitak C, Fagen R, Wilde H. Fluorescent Antibody Test for Rabies: Prospective Study of 8,987 Brains. Clin Infect Dis. 1997 Dec 1;25(6):1459–61.

9. Blenden DC, Bell JF, Tsao AT, Umoh JU. Immunofluorescent examination of the skin of rabies-infected animals as a means of early detection of rabies virus antigen. J Clin Microbiol. 1983 Sep 1;18(3):631–6.

10. Wacharapluesadee S, Tepsumethanon V, Supavonwong P, Kaewpom T, Intarut N, Hemachudha T. Detection of rabies viral RNA by TaqMan real-time RT-PCR using non-neural specimens from dogs infected with rabies virus. J Virol Methods. 2012 Sep 1;184(1):109–12.

11. Bansal K, Singh CK, Sandhu BS, Sood NK, Dandale M. Antemortem diagnosis of rabies from skin by TaqMan real time PCR. Indian J Anim Res. 2014;48(6):597.

12. Kaw A, Singh CK, Ramneek Deka, D, Sandhu BS, Awahan S, et al. Comparison of Hair Follicle and Skin for Ante Mortem Detection of Rabies: A Molecular Approach. Indian J Public Health Res Dev. 2014;5(3):165.

13. Mani RS, Madhusudana SN, Mahadevan A, Reddy V, Belludi AY, Shankar SK. Utility of real-time Taqman PCR for antemortem and postmortem diagnosis of human rabies. J Med Virol. 2014;86(10):1804–12.

14. Blenden DC, Creech W, Torres-Anjel MJ. Use of Immunofluorescence Examination to Detect Rabies Virus Antigen in the Skin of Humans with Clinical Encephalitis. J Infect Dis. 1986;154(4):698–701.

15. Dacheux L, Reynes J-M, Buchy P, Sivuth O, Diop BM, Rousset D, et al. A Reliable Diagnosis of Human Rabies Based on Analysis of Skin Biopsy Specimens. Clin Infect Dis. 2008 Dec 1;47(11):1410–7.

16. Blenden DC, Frost JW, Wachendörfer G, Dorsey C. Identification of Rabies Virus Antigen in the Skin of Foxes. Zentralblatt Für Veterinärmedizin Reihe B. 1980;27(9–10):698–704.

17. Singh CK, Ahmad A. Molecular approach for ante-mortem diagnosis of rabies in dogs. Indian J Med Res. 2018 May;147(5):513–6.

18. Smith WB, Blenden DC, Fuh TH, Hiler L. Diagnosis of rabies by immunofluorescent staining of frozen sections of skin. Amer Vet Med Assoc J [Internet]. 1972 [cited 2021 Aug 30]; Available from: https://scholar.google.com/scholar_lookup?title=Diagnosis+of+rabies+by+immunofluorescent+staining+of+frozen+sections+of+skin&author=Smith%2C+W.B.&publication_year=1972

